# A *Streptomyces venezuelae* Cell-Free Toolkit for Synthetic Biology

**DOI:** 10.1101/2020.11.16.384693

**Authors:** Simon J Moore, Hung-En Lai, Soo-Mei Chee, Ming Toh, Seth Coode, Patrick Capel, Christophe Corre, Emmanuel LC de los Santos, Paul S Freemont

**Affiliations:** Centre for Synthetic Biology and Innovation, Imperial College London, South Kensington Campus, Exhibition Road, London, SW7 2AZ, UK; Department Section of Structural and Synthetic Biology, Department of Infectious Disease; Imperial College London, South Kensington Campus, Exhibition Road, London, SW7 2AZ, UK; School of Biosciences, University of Kent, Canterbury, Kent CT2 7NJ, UK; Warwick Integrative Synthetic Biology Centre, School of Life Sciences, University of Warwick, Gibbet Hill Road, Coventry, CV4 7AL, UK; The London Biofoundry, Imperial College Translation & Innovation Hub, White City Campus, 80 Wood Lane, London W12 0BZ, UK; UK Dementia Research Institute Care Research and Technology Centre, Imperial College London, Hammersmith Campus, Du Cane Road, London, W12 0N

**Author notes:** Joint corresponding authors: Dr Simon Moore and Professor Paul Freemont.

**Keywords:** Cell-free synthetic biology, *Streptomyces*, natural products, *in vitro* transcription-translation, cell-free protein synthesis

## Abstract

Prokaryotic cell-free coupled transcription-translation (TX-TL) systems are emerging as a powerful tool to examine natural product biosynthetic pathways in a test-tube. The key advantages of this approach are the reduced experimental timescales and controlled reaction conditions. In order to realise this potential, specialised cell-free systems in organisms enriched for biosynthetic gene clusters, with strong protein production and well-characterised synthetic biology tools, is essential. The *Streptomyces* genus is a major source of natural products. To study enzymes and pathways from *Streptomyces*, we originally developed a homologous *Streptomyces* cell-free system to provide a native protein folding environment, a high G+C (%) tRNA pool and an active background metabolism. However, our initial yields were low (36 μg/mL) and showed a high level of batch-to-batch variation. Here, we present an updated high-yield and robust *Streptomyces* TX-TL protocol, reaching up to yields of 266 μg/mL of expressed recombinant protein. To complement this, we rapidly characterise a range of DNA parts with different reporters, express high G+C (%) biosynthetic genes and demonstrate an initial proof of concept for combined transcription, translation and biosynthesis of *Streptomyces* metabolic pathways in a single ‘one-pot’ reaction.

## Introduction

*Streptomyces* bacteria are environmental specialists (e.g. soil, marine, desert) that synthesize rich repertoires of natural products such as antibiotics. Much of this genetic information is locked up and cryptically regulated within biosynthetic gene clusters; regions of genomic DNA that harbor enzymes and other proteins (e.g. transporters, resistance markers). The key limitation in awakening these clusters for natural product discovery, is silent gene expression and recalcitrant genetics. Traditional strategies to overcome this include genetic modification of the host organism to bypass native regulatory elements, or the ‘capture’ of the cluster and expression in a heterologous host (1). But this can take several weeks to months to complete with varying levels of success: some cryptic clusters remain dormant due to obscure native regulation. Fundamental tools that aid these efforts, are of major interest to the natural product community.

Prokaryotic cell-free coupled transcription-translation systems are emerging as a new tool for studying natural product biosynthesis (2–8). Cell-free transcription-translation uses a crude cell-extract or purified ribosomes and translation factors – the PURE system - in a ‘one-pot’ reaction (9, 10). *E. coli* cell-extracts, referred to as either TX-TL (5, 11, 12) or cell-free protein synthesis (CFPS) (4, 13), are low-cost, straightforward to prepare and provide high recombinant protein yields, up to 2300 μg/mL (14). Moreover, metabolism is active, (15) providing ATP regeneration, while amino acid pathways are dynamic; providing additional ATP (through L-glutamate), while some amino acids deplete and are limiting for protein synthesis (16). In summary, TX-TL provides distinct opportunities for natural product biosynthesis: precursors for biosynthesis, direct control (e.g. feed precursors), short experimental timescales (4-24 hours), and stable yields. Moreover, we have shown the potential for automation in TX-TL by screening up to 500 plasmid variants in 24 hours (17).

While *E. coli* TX-TL and the PURE system are promising for natural product biosynthesis (2–4), *E. coli* has limited potential for studying biosynthetic gene clusters from *Streptomyces*, due to a number of genetic and metabolic differences: The codon content between *Streptomyces* (~70% G+C) and *E. coli* (51% G+C) is different; the regulatory sequences that control transcription, post-transcription and overall gene expression is distinct; and secondary metabolism in *E. coli* is not necessarily well-suited, or requires further metabolic engineering. Notwithstanding, *E. coli* synthesis of heterologous proteins can result in poor expression and solubility (5, 18). Therefore, we anticipate that a dedicated *Streptomyces* TX-TL system for homologous protein synthesis, has a number of advantages for studying natural product biosynthesis. As a first step, we originally released a *Streptomyces venezuelae* DSM-40230 (ATCC 10712) TX-TL system, but this had low protein yields (36 μg/mL) and showed high batch-to-batch variability. We chose *S. venezuelae* ATCC 10712, since it is well-suited to synthetic biology. *S. venezuelae* ATCC 10712 is fast-growing (40 min doubling time) and grows dispersedly in liquid culture; most *Streptomyces spp.* have slower doubling times and clump in mycelial aggregates. Moreover, *S. venezuelae* has a range of synthetic biology tools (19, 20) and is an attractive host for industrial biotechnology (21, 22). In parallel to our studies, the Jewett group (7) also established a *Streptomyces lividans* CFPS system (yields ~50 μg/mL), which was further optimised (23). A recent update to this system, highlighted the need for adding individual purified translation factors (8) to elevate protein synthesis up to ~400 μg/mL.

Based on this, we rationalized that protein synthesis, in our original proof of concept *S. venezuelae* TX-TL system, could be limited by the use of an energy solution originally optimized for a *E. coli* TX-TL protocol (11). TX-TL requires a cell-extract, a primary and secondary energy source, amino acids, cofactors, molecular crowding agents, additives (e.g. Mg^2+^, spermidine, folinic acid, tRNA) to direct protein synthesis from a template DNA sequence. The energy source is composed of nucleotide triphosphates to drive initial mRNA and protein synthesis (primary energy source) and commonly 3-phosphoglyceric acid (3-PGA) or phosphoenolpyruvate (PEP) as the secondary energy source. 3-PGA or PEP provide ATP regeneration to leverage extended protein synthesis. Potentially primary metabolism could be activated in TX-TL to provide reducing equivalents (e.g. NADH, FADH), extra energy and building blocks (e.g. amino acids, malonyl-CoA), for natural product biosynthetic pathways, as shown in cell-extract metabolic engineering (24). In this work, we focus on upgrading our *S. venezuelae* system, to elevate protein synthesis and demonstrate its broader potential for cell-free synthetic biology – for characterising DNA parts and activating some model biosynthetic pathways. To achieve this, we made some simple modifications to the system, allowing yield up to 266 μg/mL of expressed recombinant proteins and demonstrate combined transcription-translation and biosynthesis of some example natural product pathways, namely melanin and haem biosynthesis. We also describe an easy-to-follow protocol that simply requires three components: DNA, cell-extract and a master mix that we describe in detail. We believe this generic *Streptomyces* TX-TL toolkit will be of broad interest to the natural product community, complementing experimental wet-lab tools for genome mining studies.

## Results and discussion

### A high-yield *Streptomyces* TX-TL protocol

To provide an improved *Streptomyces* TX-TL toolkit for synthesis of high G+C (%) genes and pathways from *Streptomyces* spp. and related genomes, a key priority was to optimise protein production and provide a straightforward protocol with minimal batch variation, for ease of repeatability. Since bacterial transcription and translation is coupled, either these steps, physical parameters or components from the energy solution, limit overall TX-TL activity. Therefore, to keep our protocol streamlined, we made the following changes: promoter strength, energy solution, ATP regeneration and RNAse inhibition – in doing so, we obtained a high-yield protocol with minimal variation between different cell-extract batches (Figure 1A).

**Figure 1.**
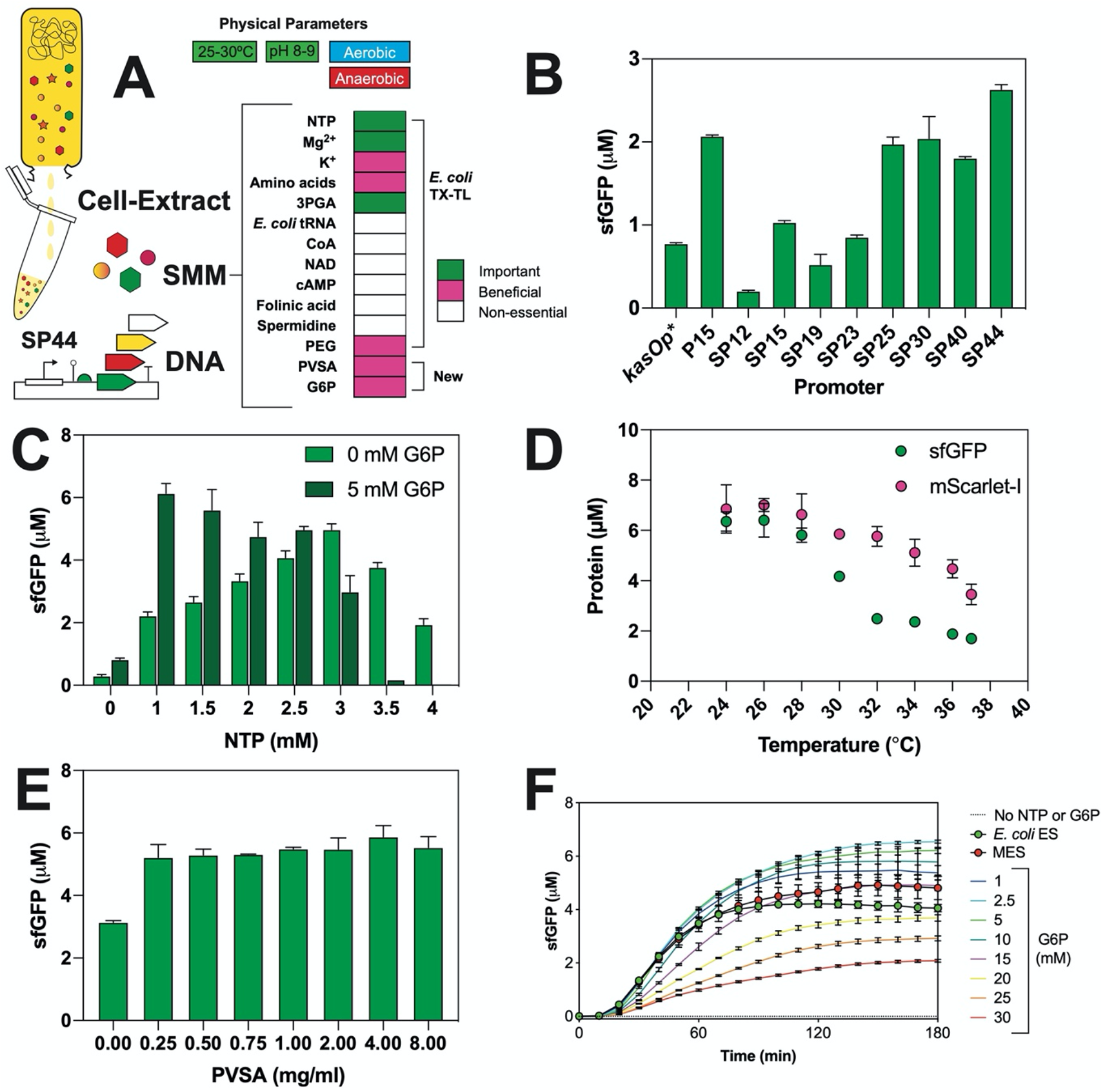
Overview of *Streptomyces* TX-TL optimisation. (A) Outline of physical and biochemical parameters of *Streptomyces* TX-TL and SMM buffer system. Optimisation of: (B) Promoter strength from Bai *et al* (19); (C) primary energy source; (D) Temperature; (E) PVSA; and (F) G6P.

#### Promoter strength

Previously, we used the *kasOp\** promoter to drive mRNA synthesis in *Streptomyces* TX-TL. This made up to 1.3 μM (36 μg/mL) of the model superfolder green fluorescence protein (sfGFP) in our previous work (5). Promoter strength is a key limiting factor in heterologous expression systems. *kasOp\** is a strong *Streptomyces* constitutive promoter, originally derived from the *kasO*/*cpkO*/*sco6280* promoter, with core −35 and −10 boxes of TTGACN and TAGART, respectively (25). The *kasOp\** promoter is active in a range of *Streptomyces* spp. through the endogenous RNA polymerases and HrdB house-keeping Sigma factor (25). Bai *et al* developed a synthetic promoter library based around *kasOp\** using fluorescence-activated cell sorting (FACS) to quantity *S. venezuelae* protoplasts (19). This included the isolation of a synthetic promoter 44 (SP44), which is 1.87-fold stronger than *kasOp\** (19). We used *Streptomyces* TX-TL to test a panel of promoters developed by Bai *et al*, with SP44 the strongest (2.63 μM sfGFP) and 2.2-fold more active than *kasOp\** (Figure 1B). We also repeated this across four independent cell-extract batches, but still observed strong batch variation. However, SP44 provided a stronger reporter plasmid to continue the optimisation process.

#### Energy solution

Next, we focused on developing a minimal energy solution (MES) to identify any non-essential components. The standard *E. coli* TX-TL energy solution used previously (5), is composed of HEPES buffer, ions (e.g. Mg-glutamate, K-glutamate), nucleotide triphosphates (NTPs - ATP, GTP, CTP and UTP), secondary energy source [typically 3-phosphoglyceric acid (3-PGA) or phosphoenolpyruvate (PEP)], amino acids, molecular crowding agent and a number of additives (13). To establish a MES for *Streptomyces* TX-TL, we first eliminated a number of non-essential components from the energy solution. This included coenzyme A, tRNA (*E. coli*), NAD, cAMP, folinic acid and spermidine (Figure S1A). While we did initially observe a positive response with cAMP, after several repeats in batches, this effect was not repeatable. For the HEPES buffer component, this was non-inhibitory (10-100 mM) and provides optimum activity between pH 8-9 (Figure S1B). For the secondary energy source, we found 3-PGA was essential; its removal decreased sfGFP synthesis by 98% (Figure S1C). We tried to replace 3-PGA with alternative secondary energy sources but observed only minimal activity: maltose (0.13 μM), sucrose (0.15 μM) and pyruvate (0.17 μM). Other potential sources such as glucose (with phosphate), PEP and succinate were inactive (Figure S1C). 3-PGA is the preferred secondary energy source in a range of non-model cell-extract hosts (26), due to its chemical stability and high energy potential, with an optimum concentration of 30 mM (Figure S1D). For the primary energy source (NTPs), there was some basal activity without additional NTPs but 3 mM ATP/GTP and 1.5 mM CTP/UTP provides peak activity (Figure S1E). Next, surprisingly the removal of amino acids only decreased sfGFP synthesis by 45%, with 0.5-1.5 mM amino acids providing peak activity (data not shown). For metal salt screen, we found that MgCl2, Mg-glutamate or Mg-acetate were all active (Figure S1F), while high levels of K-glutamate (150-200 mM) stimulated increased sfGFP synthesis (Figure S1F). This is possibly due to additional ATP regeneration via entry of α-ketoglutarate into the TCA cycle, as previously shown (27). Lastly, while we observed reasonable activity without PEG, 1% (w/v) PEG 6K was optimum, providing a 44% rise in activity (Figure S1G); it is however desirable to omit PEG for downstream natural product analytical purposes (e.g. LC-MS). Finally, based on these observations, we optimised our basic *Streptomyces* TX-TL MES system by individually fine-tuning the concentration of its core components (3-PGA, NTPs), while leaving DNA (40 nM), Mg-glutamate (4 mM), K-glutamate (150 mM), amino acids (1.25 mM) and PEG 6K (1%) constant. 3-PGA was most optimum at 30 mM, while the NTP level (ratio of 2:1 ATP/GTP:CTP/UTP), showed biphasic activity, peaking at 3 mM ATP/GTP, with full inhibition at 4 mM. Specific data on Mg-glutamate and K-glutamate optimisation with four different cell-extract batches is presented in Figure S2. As a combined result of this optimisation process, sfGFP synthesis was increased to 4 μM representing a 52% increase.

#### Additional ATP regeneration pathways

In a previous study, Caschera *et al* highlighted that other glycolytic enzymes function in *E. coli* TX-TL, using the disaccharide maltose (or maltodextrin) combined with 3-PGA to prolong protein synthesis up to 10 hours, through inorganic phosphate recycling (14). We investigated whether this part of metabolism is functional in *Streptomyces* TX-TL. Therefore, we tested the *Streptomyces* MES system (with 3-PGA) with maltose, glucose, glucose-6-phosphate (G6P) or fructose-1,6-phosphate (F16P). Interestingly, maltose, glucose, Glc6P, and F16P all prolonged the length of *Streptomyces* TX-TL activity from 2 to 3 hours. This was maximal with 5 mM Glc6P and 30 mM 3-PGA (Figure 1C), at an optimum temperature of 26°C (Figure 1D). All together we observed a 59% increase in sfGFP synthesis to 6.37 μM, but lower levels of NTP are required (Figure 1C) – equivalent to 1 mM ATP/GTP and 0.5 mM CTP/UTP. We speculate this could be related to ATP regulation of the glycolytic enzymes (e.g. hexokinase, fructokinase), leading to rapid depletion of ATP and inhibition of protein synthesis. Although this requires further investigation – there is limited literature on specific glycolytic enzymes from *Streptomyces*.

#### RNAse inhibition

As a final addition to the system, we tested the inexpensive RNAse inhibitor, polyvinylsulphonic acid (PVSA). Recently, PVSA, an RNA-mimetic, was shown to improve mRNA stability in *E. coli* TX-TL, but not increase protein synthesis (28). In *Streptomyces* TX-TL, 1 mg/mL PVSA increased sfGFP synthesis up to 5.87 μM, in the basic MES system (Figure 1E). But while we observed individual improvements with either the PVSA RNase inhibitor or the blended G6P/3-PGA secondary energy source, in combination, there was no significant additive effect with PVSA and G6P/3-PGA together. This suggested that other rate-limiting factors are at play.

In summary, we have made a specific energy solution for *Streptomyces* TX-TL with an overall 6-fold improvement in the system, partly attributed to combined use of 3PGA and G6P as secondary energy sources (Figure 1F). Furthermore, we find this can combine with remaining reaction components into a single *Streptomyces* master mix (SMM) solution, further streamlining the reaction process. With this simple modification, the TX-TL reaction requires three single components that minimises batch variation: SMM solution, plasmid DNA and the cell-extract. Next, we sought to demonstrate the use of this simplified system for the testing of plasmid tools and regulatory elements for *Streptomyces* synthetic biology.

### Cell-free characterisation of *Streptomyces* genetic tools for synthetic biology

It is highly desirable to characterise standard DNA parts using rapid and iterative design-build-test-learn cycles - the central paradigm of synthetic biology. For *Streptomyces* and related strains, either conjugation or protoplast transformation is typically used to transfer self-replicating and integrative plasmids for the testing of DNA parts for *Streptomyces* synthetic biology (19, 20, 25). DNA parts are small modular regulatory elements (e.g. promoter, insulator, tags, RBS, ORF, terminator) that facilitate downstream combinatorial DNA assembly workflows (e.g. Golden Gate) for refactoring gene expression pathways. While there are different approaches to quantitate gene expression (20, 29), Bai *et al* (19) recently applied a lysozyme method, to study single-cell gene expression quantitation of *S. venezuelae* ATCC 10712 protoplasts using fluorescence-activated cell sorting (FACS).

We next tested the promoter and RBS elements from Bai *et al* (19) in *Streptomyces* TX-TL, as well as other important regulatory elements: alternative start codons (29) and terminators (30). Firstly, we built these DNA parts to be compatible with our previous DNA assembly method EcoFlex (33). For this we had to modify the promoter consensus (prefix renamed, e.g. SP44a instead of SP44), to remove an internal BsmBI site to permit MoClo assembly. In addition, to provide comparative *in vivo* data, we built a new destination vector (cured of BsmBI and BsaI sites) from pAV-*gapdh* from Phelan *et al* (20) and renamed this StrepFlex (pSF1). pAV-*gapdh* is an integrative shuttle vector, developed as a synthetic biology plasmid tool for *S. venezuelae* (20). Firstly, for the promoter library (*kasOp\**, SP15a, SP19a, SP23a, SP25a, SP30a, SP40a, SP44a and *ermEp\**), we assembled this with the RiboJ insulator, R15 capsid RBS, mScarlet-I and the Bba_B0015 terminator. For the RBS library, *kasOp\** was used for the promoter. For the promoter variants, activity ranged from 5% (*ermEp\**) to 100% (*kasOp\**). In contrast to earlier results (Figure 1B), the BsmBI cured promoters were about 30-50% less active across the library. For the activity of the RBS variants, this ranged from 0.7% (SR9) to 117% (SR39) activity relative to the R15 capsid RBS (Figure 2B). We also tested two-dimensional promoter and RBS space with sfGFP (Figure 2C). Lastly, to provide *in vivo* data, we characterised the mScarlet-I promoter and RBS plasmids (from Figure 2A and 2B) in *S. venezuelae* ATCC 10712 (Figure S3), following the approach by Phelan *et al* (20). Interestingly, there were some significant outliers particularly for the RBS library; SR39 (along with the *E. coli* PET-RBS) was the strongest RBS in contrast to SR40, which was unexpectedly weaker both *in vitro* and *in vivo*. In addition, SR4, an expected weak RBS, was strong in both *in vitro* and *in vivo* measurements (Figure 2A-2B, Figure S3). This may reflect differences in the upstream 5’-untranslated (5’-UTR) region and the use of a different fluorescence reporter (mScarlet-I), in comparison to Bai *et al* (19). However, on the whole the promoter and RBS strengths characterised were broadly consistent with the original publication (19).

**Figure 2.**
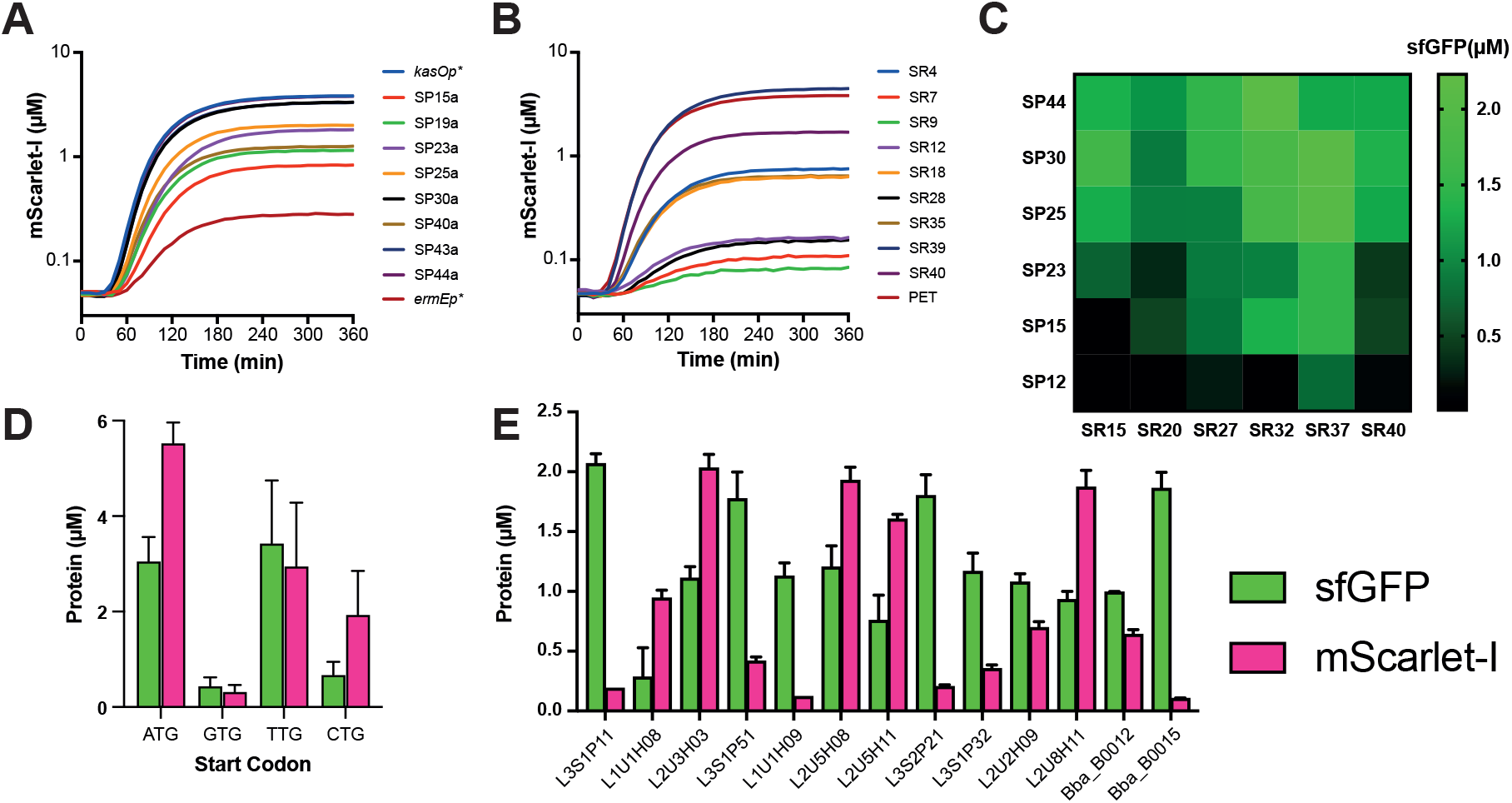
Part characterisation of *Streptomyces* regulatory elements: (A) Promoter-R15 capsid RBS-mScarlet-I; (B) *kasOp\**-RBS-mScarlet-I; (C) Promoter-RBS-sfGFP combinations; (D) variable start codons (with sfGFP and mScarlet-I); and (E) variable Rho-independent terminators from Chen *et al* (34). For terminator plasmid design, see Moore *et al* (33). 40 nM plasmid DNA was incubated in the optimised reaction conditions at 28°C as a technical triplicate repeat and repeated on two separate days. Unless otherwise stated, the SP44 promoter, PET RBS and Bba_B0015 were used in constructs, assembled into either pTU1-A (*E. coli*) or pSF-1 (*E. coli* and *Streptomyces* shuttle vector).

Another important, but perhaps underused regulator of gene expression in the context of synthetic biology, is the start codon. In most bacteria, ATG is the preferred codon for translation initiation through fMet-tRNA. Previously, Myronovskyi *et al* used a β-glucuronidase (GUS) reporter to show that the TTG codon was stronger than ATG for translation initiation, by almost 2-fold in both *Streptomyces albus* J1074 and *Streptomyces sp.* Tu6071 (29). Using sfGFP and mScarlet-I as reporters, our findings suggest that for *S. venezuelae* at least, ATG is equivalent in strength to TTG, followed by CTG and lastly GTG as the weakest (Figure 2C). This also likely changes with coding sequences and 5’-UTR. In comparison, for *E. coli* the order of strength goes as follows: ATG > GTG > TTG > CTG (31). We expect this differs due to high GC codon bias in *Streptomyces*. Despite the use of different experimental conditions, our results confirm that TTG is a strong alternative start codon and that GTG is weak, although the role of this is unclear and intriguing due its high frequency in *Streptomyces* genomes (32).

Lastly, to the best of our knowledge, no studies have so far reported the use of terminators for controlling pathway expression in *Streptomyces*. Using the same experimental format as we previously used in EcoFlex (33), we tested a selection of Rho-independent terminators from the iGEM catalogue (Bba_B0012, Bba_B0015) and from Chen *et al* (34) in *S. venezuelae* TX-TL (Figure 2D). These activities strongly follow our previous observations in *E. coli* cell-free (33) and these terminators provide a final tool for the design and assembly of synthetic metabolic pathways. These terminators were also designed to prevent repetition in DNA elements and protect against homologous recombination as previously highlighted (34, 35). For now, our TX-TL system demonstrates proof-of-concept data for prototyping DNA parts in *Streptomyces*, which may in future provide a platform for designing pathways – for refactoring precise synthetic biology control of metabolic pathways in engineered *Streptomyces* strains.

### TX-TL synthesis of high G+C (%) genes

Previously, we found our *Streptomyces* TX-TL system was most active with 40 nM of plasmid DNA, using the *kasOp\**-sfGFP reporter, to saturate protein synthesis (28). In comparison to *E. coli* TX-TL, protein synthesis is saturated at around 5-10 nM of reporter DNA, which varies with different promoters and sigma factors (36). We questioned whether DNA degradation led to this discrepancy; most *Streptomyces spp.* degrade methylated plasmid DNA with endonucleases (37). To compare methylated and unmethylated plasmid DNA for methylation-specific endonucleases, we tested unmethylated and methylated SP44-sfGFP reporter plasmid in *Streptomyces* TX-TL. Interestingly, there was no major change in sfGFP synthesis between unmethylated or methylated plasmid DNA, across different DNA concentrations (Figure S4). We also tested relative plasmid DNA stability in *S. venezuelae* cell-extracts, with the standard MES energy solution, and incubated at different time-lengths, followed by re-extraction of the plasmid DNA (using the Qiagen plasmid DNA purification kit), which was then separated and visualised on a 1% (w/v) agarose gel. This indicated that methylated plasmid DNA is stable, during the time (0-4 hr) when TX-TL is active (Figure S4). Further to this, we also tested linear DNA for exonuclease activity. To protect the coding sequences, we PCR amplified about 150-250 bases upstream and downstream of the coding parts, using the standard SP44-sfGFP reporter plasmid. But in the TX-TL reaction, linear DNA was 95% less active than circular DNA, at 40 nM of DNA (Figure S5). This suggests the *S. venezuelae* cell-extract has exonuclease activity, but endonuclease activity is minimal.

Since circular DNA degradation was not a limiting factor, we tested different fluorescent proteins (Figure 3A) to determine if the optimum plasmid DNA concentration, for protein synthesis, changes. Firstly, we tested mVenus-I and mScarlet-I, combined with the strong SP44 promoter, in comparison to the SP44-sfGFP reporter. The maximum yields achieved for these three proteins were: 6.48 μM sfGFP (174 μg/mL), 9.50 μM mScarlet-I (266 μg/mL) and 7.72 μM mVenus-I (224 μg/mL). Both SP44-mScarlet-I and SP44-mVenus saturated protein synthesis with a lower DNA template (10 nM) (Figure 3B). This was surprising, since the coding sequence of mVenus is 96% identical to sfGFP, with the exception of 30 mutations and an additional GTG (valine) at the second codon for mVenus-I. We speculate that this alters either mRNA stability or translation initiation, although this requires further investigation. Secondly, we also tested the robustness of the system for other proteins from high G+C (%) genes (Figure 3C), using the oxytetracycline enzymes (OxyA, -B, -C, -D, -J, -K, -N and -T) from *Streptomyces rimosus* that were previously only detectable by Western blotting in our original publication (30), as well as three non-ribosomal peptide synthetases (NRPS); this included the TxtA and TxtB NRPS enzymes from thaxtomin A biosynthesis in *Streptomyces scabiei* and an uncharacterised NRPS (NH08_RS0107360) from *S. rimosus*. With the exception of TxtA, most enzymes were discernible by either SDS-PAGE (Figure 3C), or for OxyA (47 kDa) and TxtB (162 kDa), low levels (<0.5 μM) were detected by Western blotting using an anti-Flag tag (data not shown). We also incorporated a C-terminal tetracysteine tag with the oxytetracycline enzymes and mScarlet-I, using the fluorogenic biarsenical dye fluorescein arsenical hairpin binder-ethanedithiol (FlAsH-EDT_2_), to measure real-time nascent protein synthesis (Figure 3D). Most oxytetracycline enzymes showed a significant increase in FlAsH-EDT_2_ fluorescence (P < 0.05), peaking at 120 min, with only OxyN producing the weakest response (P = 0.056), although clearly detectable by PAGE or Western. For mScarlet-I, the time-lag between the fluorescence signals for FlAsH-EDT2 (immature protein) and mScarlet-I (mature protein), allowed us to estimate a maturation time of 40 min for mScarlet-I (Figure 3E); this is in close agreement to a literature value of 36 min, calculated *in vivo* (38). In summary, our *Streptomyces* TX-TL system is robust for expression of high G+C (%) genes, has multiple tools (e.g. tags, plasmid systems) and is comparable to other bacterial TX-TL systems.

**Figure 3.**
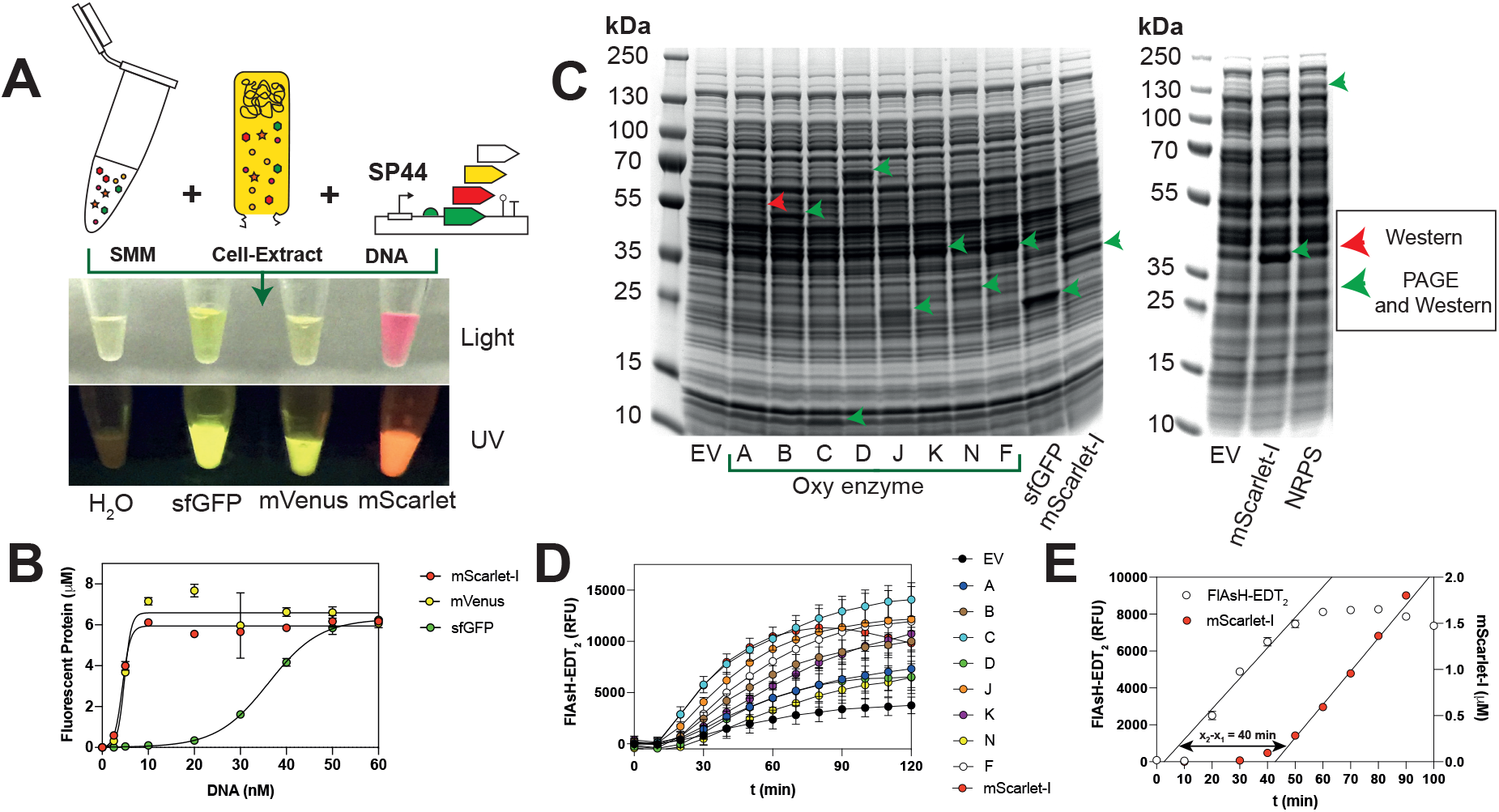
Robust and high-yield synthesis of high G+C (%) genes. (A) Synthesis of codon-optimised fluorescence proteins. (B) Denaturing PAGE of oxytetracycline biosynthetic proteins, fluorescence proteins and a representative NRPS (NH08_RS0107360 from *S. rimosus*). (C) Saturation of protein synthesis for sfGFP, mVenus and mScarlet-I with increasing DNA concentrations. (D) Real-time detection of protein synthesis with C-terminal FlAsH-EDT_2_ tag system. (E) Estimation of mScarlet-I maturation time with real-time measurement of immature and mature protein synthesis.

### Transcription, translation and biosynthesis

The next step was to reconstitute a biosynthetic pathway in *Streptomyces* TX-TL system. Initially, to show the synthesis of a single enzyme and its activity, we selected the GUS reporter enzyme. We synthesised the enzyme in the TX-TL reaction from 40 nM SP44-*gus*, left for 4 hrs at 30°C. The GUS enzyme showed a clear band on SDS-PAGE at the expected size of 68 kDa (Figure S6). To test for GUS activity, an equal volume of the TX-TL extract, as well as a negative control reaction, was mixed with X-GlcA substrate in increasing concentrations (10, 25 and 50 mg/mL). Only the extract from the SP44-*gus* reaction developed a deep blue pigment, within minutes, indicating strong GUS activity (Figure 4A).

**Figure 4.**
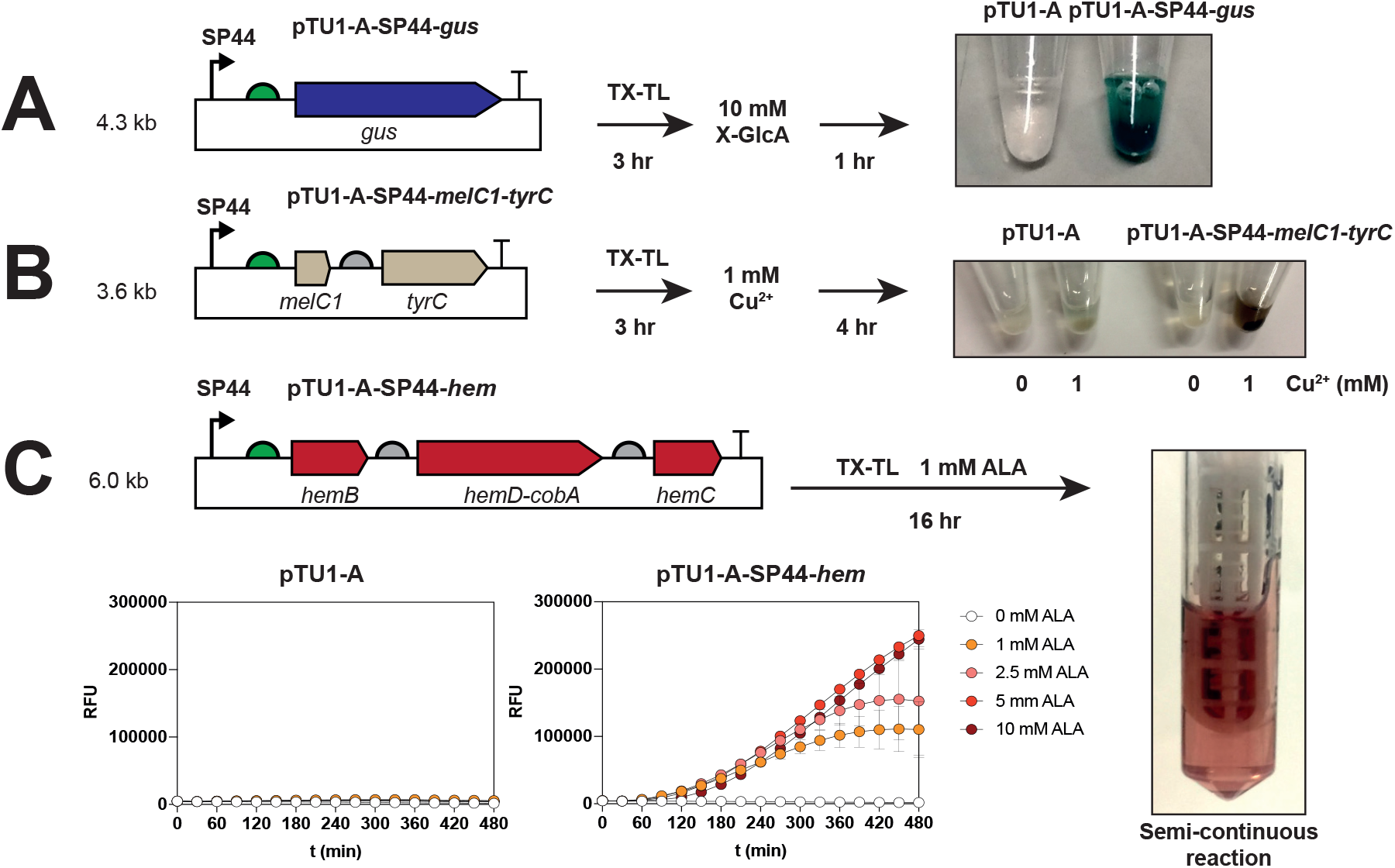
*Streptomyces* cell-free transcription, translation and biosynthesis. (A) Codon-optimised *E. coli* MG1655 GUS enzyme. (B) *S. venezuelae* DSM-40230 tyrosinase (TyrC) and copper metallochaperone (MelC1). (C) *S. venezuelae* DSM-40230 early-stage haem biosynthetic pathway, HemB, HemD-CysG^A^ and HemC. Addition of individual substrates and approximate timescales are indicated within the diagram. For details of batch and semi-continuous reaction conditions, please refer to methods.

Next, we selected two metabolic pathways to provide a further test for the TX-TL system. We selected two operons from *S. venezuelae* encoding the melanin and early-stage haem biosynthetic pathways to provide a discernible output for testing (fluorescence and or colorimetric). Also, both operons were selected from *S. venezuelae*, to improve expression in TX-TL, since the codon usage is adapted to this host.

Melanin is a natural pigment that absorbs ultraviolet (UV) light to protect cells from DNA damage. Recently, Matoba *et al* studied the mechanism of tyrosinase and the role of the “caddie” protein from *Streptomyces castaneoglobisporus* HUT6202 (39). Tyrosinase, TyrC, catalyses the rate-limiting step in melanin biosynthesis: it oxidises the phenol group (in L-tyrosine) into the *orthro*-quinone intermediate, which enters an autocatalytic cascade into the melanin pathway. TyrC is dicopper-dependent, with each Cu(II) atom coordinated by three His residues, which is transferred by MelC1, a small (12.8 kDa) metallochaperone, which transfers copper to the active site. The *S. venezuelae* tyrosinase operon encodes both MelC1 and TyrC. Using our *Streptomyces* TX-TL and SP44-*melC1*-*tyrC*, from denaturing PAGE, we saw clear synthesis of TyrC at approximately 34 kDa (expected 31.4 kDa) although MelC1 was indistinguishable (Figure S7). In terms of activity, we observed brown pigment formation after ~2 hours, only with the addition of 1 mM CuCl_2_ (Figure 4B). This indicated melanin biosynthesis was active, despite the apparent absence of MelC1. Without the addition of CuCl_2_, or the tyrosinase plasmid, the cell-extracts remained clear. Previously, Matoba *et al* showed insertion of Cu(II) into TyrC by MelC1 involves a transient interaction, and that MelC1 is unstable and forms aggregates that are difficult to detect with PAGE (39). Also, apo-TyrC is inactive with Cu(II) alone, which suggests that our TX-TL system supports the synthesis of both TyrC and MelC1.

Lastly, we tested a three-gene biosynthetic operon (*hemC*-*hemD*/*cysG^A^-hemB*) that catalyses the early-stages of haem biosynthesis (40). This pathway was selected since it contains a known fluorescence reporter enzyme CysG^A^ (41) (also referred to CobA), a methyltransferase naturally fused as HemD/CysG^A^. We added a pTU1-A-SP44-*hemC*-*hemD*/*cysG^A^*-*hemB* (pTU1-A-SP44-*hem*) plasmid into the TX-TL reaction, as well as a negative control plasmid (pTU1-A), both with and without 5-aminolevulinic acid (5-ALA), the substrate for the pathway. In the presence of pTU1-A and 1 mM ALA, there was some minor background fluorescence (Figure 4C), which we expected since haem biosynthesis is essential. In contrast, with pTU1-A-SP44-*hem* and 1 mM ALA, this generated strong red fluorescence, 20-fold higher than background levels in the control reaction (pTU1-A and 1 mM ALA). For protein synthesis, while we could detect HemB (35 kDa) and HemC (38 kDa), the fusion protein HemD/CysG^A^ was less clear, with other major bands at the expected mass – 58.3 kDa (Figure S6). To verify pathway function, we ran a semi-continuous reaction (42) to facilitate purification by separating the haem intermediates from the cell-extract proteins (inset image in Figure 4C). Interestingly, LC-MS analysis detected the air-oxidised product of the HemD enzyme (uroporphyrinogen III - 837 m/z), observed as a 6-electron oxidised uroporphyrin III (red fluorescent) intermediate at 831 *m/z*; typical for these air-sensitive intermediates (Figure S8). In an attempt to limit oxidation, we also repeated these assays under anaerobic conditions using a layer of mineral oil. Surprisingly, while the TX-TL reactions were still active (including for sfGFP/mScarlet-I), this was insufficient to prevent oxidation of uroporphyrinogen III to uroporphyrin III (data not shown). These results show that our *S. venezuelae* TX-TL system can support the synthesis of at least three enzymes from plasmid DNA in a combined ‘one-pot’ combined translation, translation and enzymatic pathway.

## Conclusions

Our study complements a recent surge in interest in the use of cell-free systems for the study of biosynthetic pathways (2, 4, 23, 42). Here we wanted to expand the palette of plasmid tools for the further development of *S. venezuelae* as a synthetic biology chassis by developing an optimised streptomyces TX-TL toolkit (5, 20, 21). Our combined findings show an order of magnitude improvement in protein synthesis over our original streptomyces TX-TL system, without the requirement of any genetic modifications to either limit RNA degradation or increase translation rates; indeed, Xu *et al* recently show this is clear rate-limiting step for other *Streptomyces* cell-free systems (8). Unexpectedly, we found the system functions anaerobically, which in the future may facilitate the study of oxygen-sensitive pathways. We also demonstrate that the semi-continuous system permits reasonable milligram scale-up of biosynthetic metabolites and a clean route to purification and analysis. In conclusion, our results realise the early-stage potential of *Streptomyces* cell-free for the study of synthetic biology for natural products, providing a native prototyping environment for developing synthetic biology tools (e.g. promoters/RBS) and also for exploring biosynthetic pathways from these organisms.

## Methods

### Molecular biology

All plasmids were either prepared using EcoFlex cloning or by routine materials and methods, as previously described (34). For PCR of high GC genes and operons, Q5 polymerase (NEB, UK) was used, using standard cycling or touchdown (72-59°C annealing) and the addition of 5% DMSO. For tricky amplicons, the protocol was modified with an annealing time of 30 seconds and elongation temperature reduced to 68°C. The following bacterial strains were used: *S. venezuelae* DSM-40230 and *E. coli* DH10β. Unmethylated plasmid DNA was prepared from an *E. coli dam*^*−*^*dcm*^*−*^ mutant (C2925) obtained from NEB. Plasmids and oligonucleotides are listed in Table S1-S5.

### Preparation of cell-extracts

*S. venezuelae* ATCC 10712 was grown in GYM (prepared in distilled water). The cell-extracts were prepared as described previously (33), with the exception that β-mercaptoethanol was removed from the wash buffers and replaced with 2 mM dithiothreitol.

### Energy solution and reaction conditions

The reaction mixture contained: 8 mg/mL cell-extract, 40 nM DNA template, 25 mM HEPES, 1 mM ATP, 1 mM GTP, 0.5 mM UTP, 0.5 mM CTP, 30 mM 3-PGA, 5 mM glucose-6-phosphate, 1.5 mM amino acids (1.25 mM L-leucine), 4 mM Mg-glutamate, 150 mM K-glutamate, 1% (w/v) PEG6K and 5 mg/mL PVSA. All reactions were incubated at 30°C, 40 nM pTU1-A-SP44-sfGFP and the MES or SMM buffer system, unless otherwise stated. At least three technical repeats were prepared (for fluorescence measurements) and repeated with at least two independent cell-extract batches (from A4-A7) prepared on separated days.

### Denaturing PAGE

40 μL cell-free reaction (30 μL SMM + 10 μL plasmid DNA) was incubated in a 2 mL tube at 25-30°C (no shaking) for 6 hours. To precipitate proteins, 1 mL ice-cold 100% (v/v) acetone was added. Samples were placed at −20°C for 30 mins, before centrifugation at 18,000 × *g*, 4°C for 10 min. The supernatant was removed, and the pellet was washed with 1 mL ice-cold 70% (v/v) acetone. Centrifugation and supernatant removal steps were then repeated. The pellet was air-dried, before re-suspending in 30 μL ddH_2_O and 10 μL 4X NuPAGE lithium dodecyl sulphate (LDS) sample buffer (ThermoFisher) and boiled at 100°C for 5 min. To ensure the pellet was solubilized, samples were aspirated with a pipette five times, and if necessary (a visible pellet remaining), left for an additional 5 min at 100°C. 10-100 μg total protein was then separated with a 4-12% (v/v) gradient Bis-Tris gel (ThermoFisher) run in MES buffer system. Proteins were stained with InstantBlue (Generon), destained with ddH_2_O and images were recorded with the ChemiDoc XRS imaging system (Biorad).

### TX-TL fluorescence measurements

10 μL cell-free reactions were prepared in a 384-well black Clear®, F-bottom, low-binding plate (Greiner). Reactions were measured as a triplicate technical repeat and at least repeated with cell-extracts prepared from two separate days. were measured in a 384-well plate. The plate was sealed with aluminium film, SILVERseal™ (Greiner), briefly centrifuged at 2,000 × *g* for 10 seconds. Real-time plate measurements were recorded in a CLARIOStar**©** plate reader (BMG Labtech, Germany) at 30°C with 10 seconds of shaking at 500 rpm prior to measurements, using either standard filters (Omega) or monochromator settings (CLARIOStar). Purified sfGFP, mVenus-I and mScarlet-I standards were purified, as described previously (33), to estimate protein concentration during real-time fluorescence measurements.

### Mass spectrometry analysis

TX-TL reactions were prepared as two components (A and B) in a semi-continuous reaction as follows: Component A - 100 μL of standard TX-TL reaction, in the absence of PEG, was injected into a Thermo Scientific Pierce 3.5K MWCO 96-well microdialysis device; Component B - 1.5 mL SMM solution with 1 mg/mL carbenicillin, in a 2.5 mL tube. The microdialysis cassette was placed inside the 2.5 mL tube and incubated at 30°C for 24 hrs with shaking (1000 rpm). Samples were acidified with 1% (v/v) HCl, centrifuged at 18,000 x *g* for 25 min at room temperature. The supernatant was loaded onto a pre-equilibrated, with 1% (v/v) HCl, Sep-Pak C-18 (50 mg sorbent) solid-phase extraction cartridge (Waters), washed with 10 mL of 10% (v/v) ethanol and eluted with 2 mL of 50% (v/v) ethanol. All solutions were acidified with 1% (v/v) HCl. Eluted samples were dried under vacuum, at room temperature, using an Eppendorf Concentrator Plus. Samples were dissolved in 150 μL 1% HCl and centrifuged again at 18,000 x *g* for 25 min at room temperature. 1 μL of supernatant was then analysed by LC-MS, performed with an Agilent 1290 Infinity system with an online diode array detector in combination with a Bruker 6500 quadruple time-of-flight (Q-ToF) mass spectrometer. An Agilent Extend-C18 2.1 x 50mm (1.8 μm particle size) column was used at a temperature of 40°C with a buffer flow rate of 0.5 mL/min. LC was performed with a gradient of buffer A (0.1% formic acid in water) and buffer B (0.1% formic acid in acetonitrile). Separation was achieved using 2% buffer B for 0.6 min, followed by a linear gradient to 100% buffer B from 0.6 - 4.6 min, which was held at 100% buffer B from 4.6 - 5.6 min followed by a return to 2% buffer B from 5.6 - 6.6 min, along with 1 min post run. Spectra were recorded between a mass range of 50-1700 *m/z* at a rate of 10 spectra per second in positive polarity.

## Acknowledgments

The authors would like to thank Professor Jay Keasling (University of California) for providing the plasmid pAV-*gapdh*.

