## Supplementary Information for "A *Streptomyces venezuelae* Cell-Free Toolkit for Synthetic Biology"

#### *S. venezuelae* ATCC 10712 promoter and RBS platereader characterisation

Cells were grown in 5 mL GYM media, 25 µg/mL apramycin as two biological repeats and three technical repeat measurements. 1 mL of cells was centrifuged 3,350 x g for 10 min, and the cell pellet was washed with 1 mL dd$H_2O$. 100 µL of culture was then measured for absorbance at 600 nm and fluorescence in a Greiner 96-well black plate (clear bottom) on a Clariostar® (BMG biotech) platereader, using the following settings: sfGFP excitation 470-15 nm, emission 515-20 nm and 1200 gain; mScarlet-I excitation 583-15 nm, emission 626-20 nm and 3000 gain.

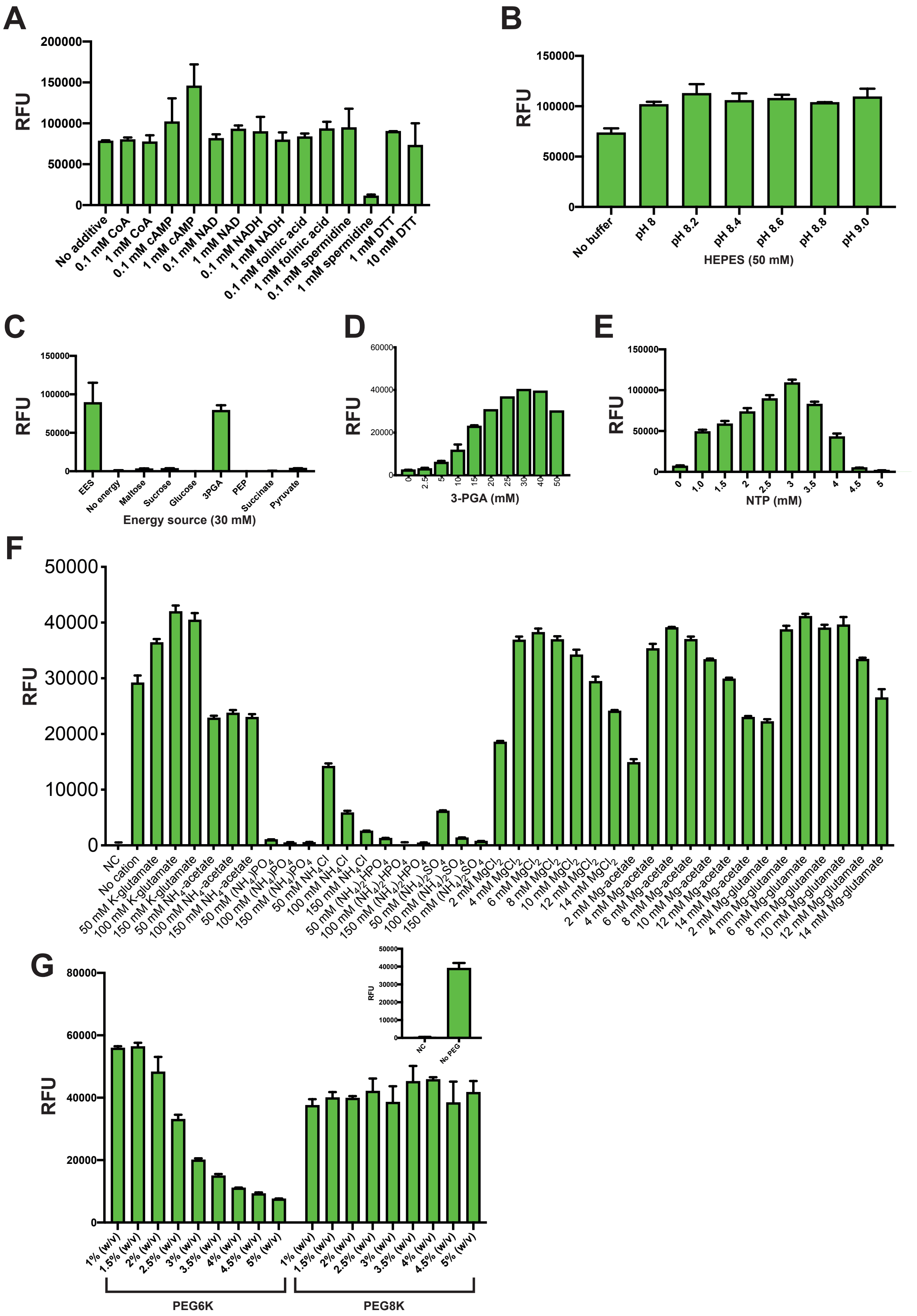


**Figure S1.** Optimisation of the MES buffer. (A) Additives, (B) 50 mM HEPES pH, (C) energy source (30 mM), (D) 3-PGA (mM), (E) NTP concentration (2:1 ratio of ATP/GTP to CTP/UTP). (F) metal salts and (G) PEG. For all experiments 40 nM pTU1-A-SP44-PET-sfGFP-Bba_B0015 was used, with standard conditions outlined in the methods. Data is representative of three technical repeats, but experiments were repeated in at least two different extract batches (biological repeat) to observe for consistent trends. The positive effect of cAMP in panel A was not repeatable under different batches. Abbreviations used within figure: *E. coli* energy solution (EES); dithiothreitol (DTT); phosphoenolpyruvate (PEP); negative control (NC).

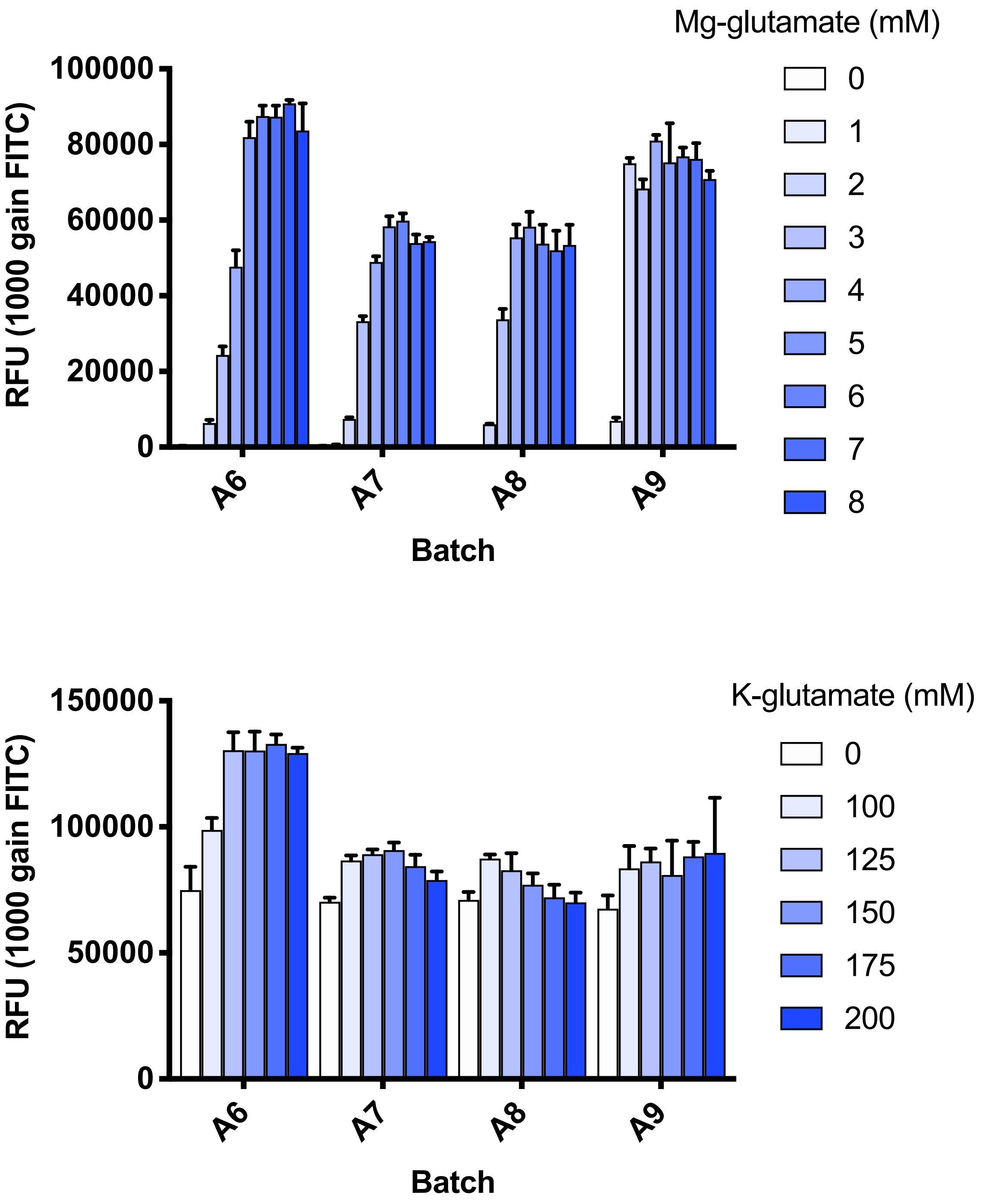


**Figure S2.** Mg-glutamate and K-glutamate (mM) optimisation with A6-A9 cell-extract batches. Experimental conditions are identical to Figure S1 using the MES buffer.

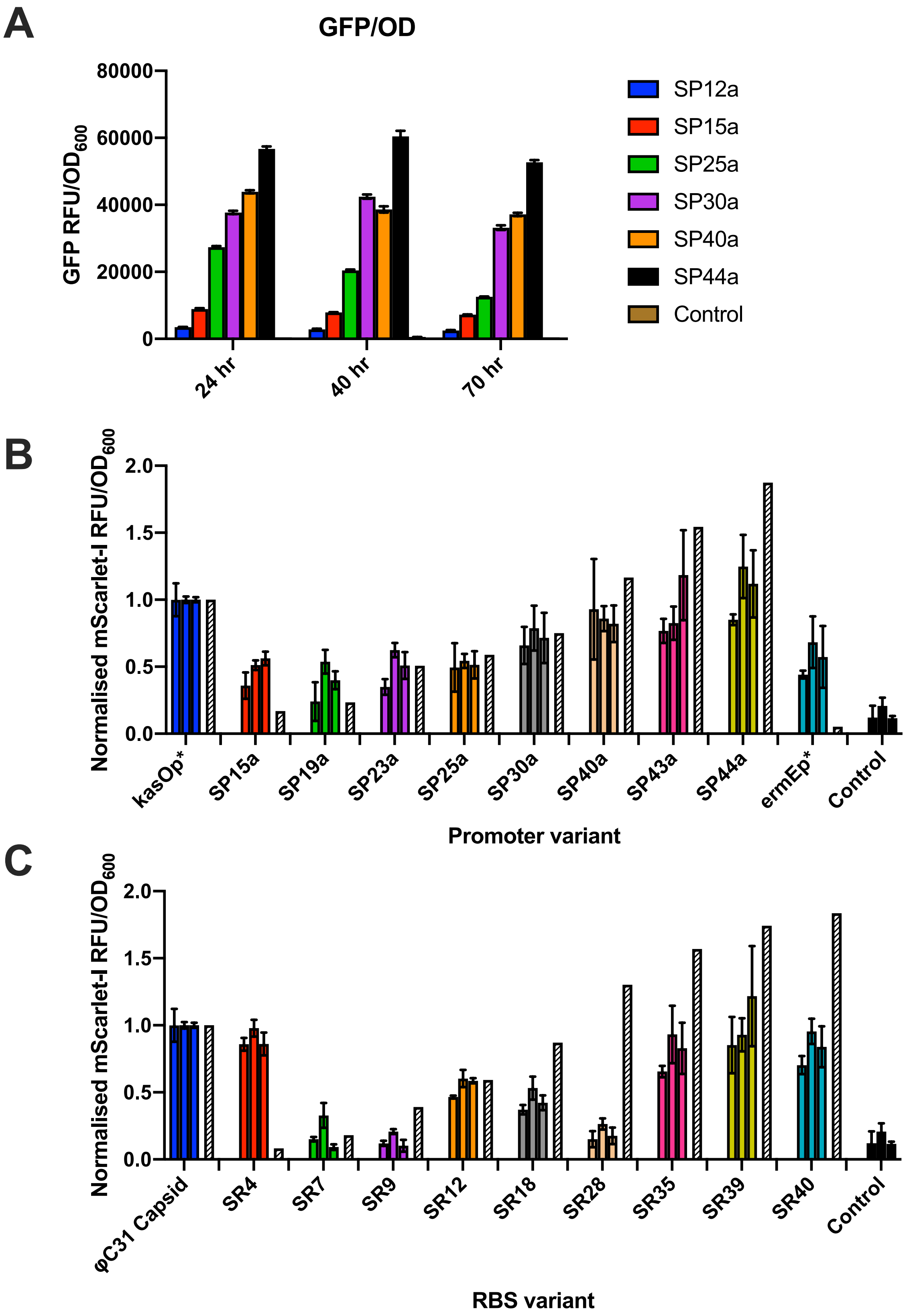


**Figure S3.** Characterisation of select promoters and RBS elements in *S. venezuelae* ATCC 10712. (A) Timepoint measurement of the pSF1-sfGFP promoter library at 24 hr, 40 hr and 70 hrs. (B) Normalised promoter strength of the pSF1-mScarlet-I library. (C) Normalised RBS strength of the pSF1-mScarlet-I library. The control culture is representative of an empty vector (pSF-1) for measurement of background fluorescence. White dashed bars represent predicted promoter/RBS strength from Bai *et al*.

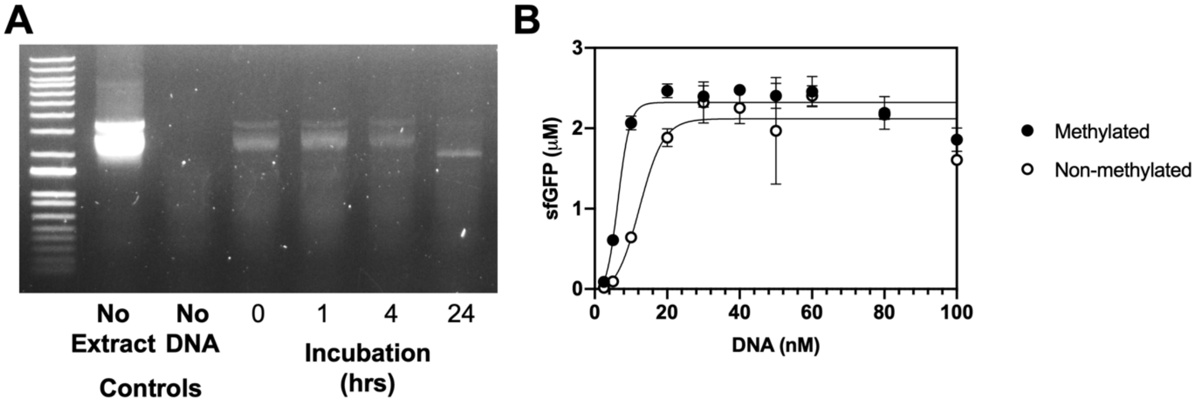


**Figure S4. Methylated DNA is not degraded in *S. venezuelae* ATCC 10712 cell-extracts during the active reaction period.** (A) 40 nM methylated pTU1-A-SP44-PET-sfGFP-Bba_B0015 was incubated in 33 μL of a standard 10 mg/mL cell-free experiment with energy solution, and purified on a mini-prep (Qiagen) column after different time points. (B) Incubation of 40 nM methylated and non-methylated pTU1-A-SP44-PET-sfGFP-Bba_B0015 in a standard *S. venezuelae* TX-TL reaction at different DNA concentrations.

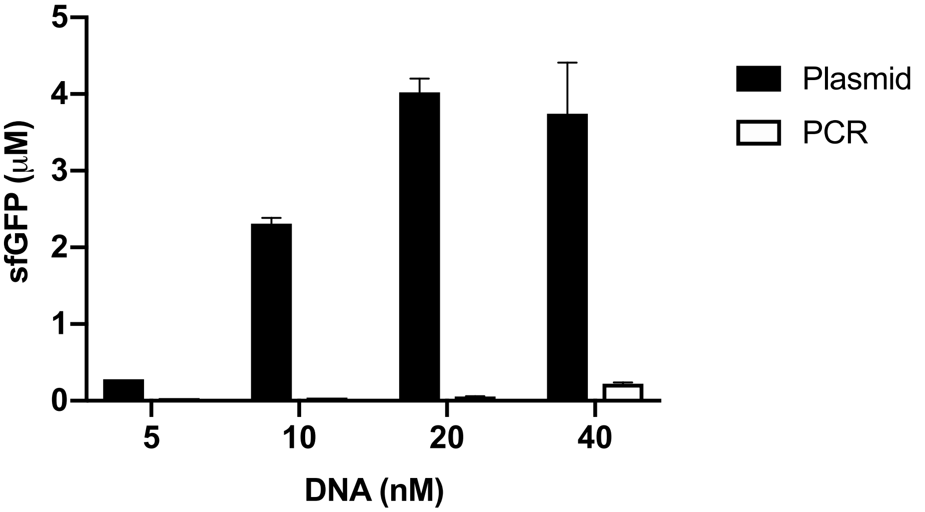


**Figure S5.** *S. venezuelae* TX-TL activity of plasmid and PCR products under optimised conditions. Plasmid reporter: pTU1-A-SP44-PET-sfGFP-Bba_B0015

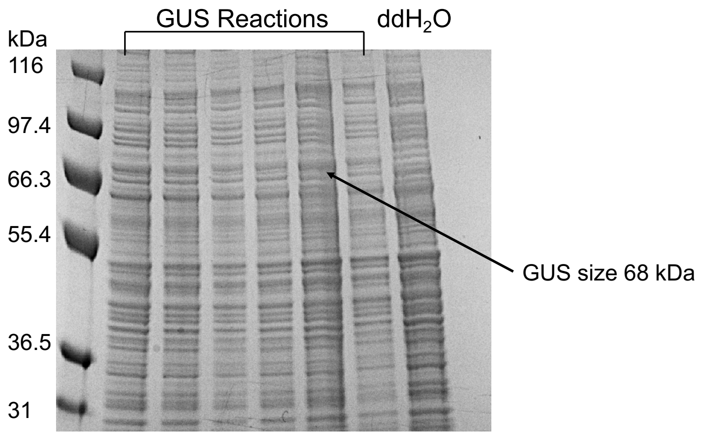


**Figure S6.** SDS-PAGE of GUS synthesis in *S. venezuelae* TX-TL

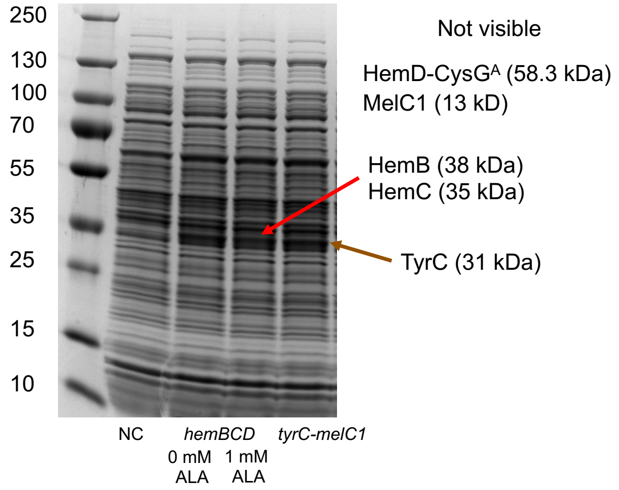


**Figure S7**. SDS-PAGE of the melanin and heam operons in *S. venezuelae* TX-TL.

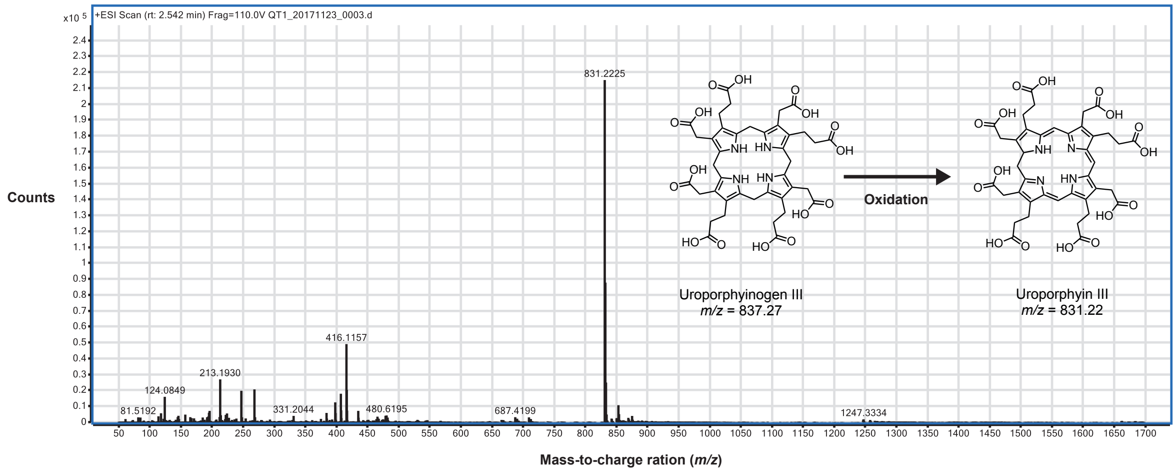


**Figure S8.** Mass spectrometry of the uroporphyrin III intermediate isolated from the semi-continuous reaction (from Figure 4). See methods for further details.

### Table S1 – Plasmids created in this study

| Plasmid ID | Description | Marker |
|---|---|---|
| pSJM899 | pBP-*kasOp** | $Cam^R$ |
| pSJM1044 | pBP-SP44 | $Cam^R$ |
| pSJM1047 | pTU1-A-RFP | $Amp^R$ |
| pSJM1059 | pTU1-A-SP44-PET-*sfGFP*-Bba_B0015 | $Amp^R$ |
| pSJM1113 | pET15b-*sfGFP* | $Amp^R$ |
| pSJM1147 | pTU1-A-SP44-PET-*hemCDB*-Bba_B0015 | $Amp^R$ |
| pSJM1174 | pTU1-A-SP44-PET-*mScarlet*-Flag-FlAsH-dBrocolli Bba_B0015 | $Amp^R$ |
| pSJM1177 | pTU1-A-SP44-PET-*melC1-tyrC*-Bba_B0015 | $Amp^R$ |
| pSJM1180 | pTU1-A-SP44-PET-*oxyA*-Flag-FlAsH-dBrocolli-Bba_B0015 | $Amp^R$ |
| pSJM1181 | pTU1-A-SP44-PET-*oxyB*-Flag-FlAsH-dBrocolli-Bba_B0015 | $Amp^R$ |
| pSJM1182 | pTU1-A-SP44-PET-*oxyC*-Flag-FlAsH-dBrocolli-Bba_B0015 | $Amp^R$ |
| pSJM1183 | pTU1-A-SP44-PET-*oxyD*-Flag-FlAsH-dBrocolli-Bba_B0015 | $Amp^R$ |
| pSJM1184 | pTU1-A-SP44-PET-*oxyJ*-Flag-FlAsH-dBrocolli-Bba_B0015 | $Amp^R$ |
| pSJM1185 | pTU1-A-SP44-PET-*oxyK*-Flag-FlAsH-dBrocolli-Bba_B0015 | $Amp^R$ |
| pSJM1186 | pTU1-A-SP44-PET-*oxyN*-Flag-FlAsH-dBrocolli-Bba_B0015 | $Amp^R$ |
| pSJM1187 | pTU1-A-SP44-PET-*oxyT*-Flag-FlAsH-dBrocolli-Bba_B0015 | $Amp^R$ |
| pSJM1190 | pTU1-A-SP44-PET-*NH08_RS0107360*-Flag-FlAsH-dBrocolli-Bba_B0015 | $Amp^R$ |
| pSJM1192 | pET15b-*mScarlet* | $Amp^R$ |
| pSJM1192 | pET15b-*mVenus* | $Amp^R$ |
| pSJM1356 | pTU1-A-SP44-PET-*txtA*-Flag-FlAsH-dBrocolli-Bba_B0015 | $Amp^R$ |
| pSJM1357 | pTU1-A-SP44-PET-*txtB*-Flag-FlAsH-dBrocolli-Bba_B0015 | $Amp^R$ |
| pSJM1379 | pSF1-2A-RFP (StrepFlex, accepts four EcoFlex Level 1 plasmids) | $Apr^R$ |
| pSJM1380 | pSF1-2a-RFP (StrepFlex, accepts two EcoFlex Level 1 plasmids) | $Apr^R$ |
| pSJM1381 | pSF1-2b-RFP (StrepFlex, accepts three EcoFlex Level 1 plasmids) | $Apr^R$ |
| pSJM1485 | pTU1-A-SP44-PET-*mVenus*-Flag-FlAsH-Bba_B0015 | $Amp^R$ |
| pSJM1486 | pTU1-A-SP44-PET-*gus*-Flag-FlAsH-Bba_B0015 | $Amp^R$ |

Abbreviations: Flag (Flag tag) – DYKDDDDK; FlAsH (FlAsH-$EDT_2$ tag) – CCPGCC

### Table S2 – PCR oligonucleotides

Colour Code

GGTCTCANNNN or NNNNAGAGACC – BsaI

CATATG – NdeI

AGATCT – BglII

GGATCC – BamHI

TGTACA – BsrGI

GAC…AAG – Flag-tag

TGC…TGC – FlAsH-tag

TTG…CAA – dBrocolli aptamer

| Oligo ID | Sequence (5'-3') | Template |
|---|---|---|
| melC1-tyrC_NdeI_F | CACCATATGTCCAACATCACCCGCCGCCGGGCCCTC | *S. venezuelae* DSM-40230 |
| melC1-tyrC_BamHI_R | CACGGATCCGGCGGCGGTGTCGAAGGTGTAGTGG | |
| hemC_NdeI_F | CACCATATGACCGAGAGGGCACTGAG | |
| hemB_BglII_R | CACAGATCTCTAGCCGCGCGAGGCGGAC | |
| oxyA_F | CACCATATGTCCAAGATCCATGACGCGCGAC | *S. rimosus* DSM-40260 |
| oxyA_R | GTGGGATCCCCGCTGCCTCCCCGGCTC | |
| oxyB_F | CACCATATGACCGGCCAGCTCGCCCCCGCT | |
| oxyB_R | CACGGATCCGTCCCGGCCGCTGACCACC | |

| | | |
|---|---|---|
| oxyC_F | CACCATATGACCCTGCTCACCCTCTCCGAC | |
| oxyC_R | GTGGGATCCCTTGTCCCGCGCGGCGCCAC | |
| oxyJ_F | CACCATATGACTACCAGTGCCACGCCCAG | |
| oxyJ_R | GTGGGATCCGTAAATGCCCAGCCCGCCGCAG | |
| oxyK_F | CACCATATGCCCGCACCCACATCCCAC | |
| oxyK_R | GTGGGATCCGGCGGCATGCGGGGCCTC | |
| oxyN_F | CACCATATGCGCATCATCGATCTGTCGACAACC | |
| oxyN_R | GTGGGATCCCTCCTCCACCACCGCCACC | |
| oxyF_F | CACCATATGACCAGCACCTCTCCCACGGCGCTTC | |
| oxyF_R | CACGGATCCCGCGGGCACCGTGCTCAC | |
| NH08_RS0107360_NdeI_F | CACCATATGACGGCCGTACACGAGATC | |
| NH08_RS0107360_BamHI_R | CACGGATCCCGGCCGGTCCGGCTGCCGCAGGAG | |
| txtA_NdeI_F | CACCATATGTCGCACCTGACCGGTGAAGATCTCC | *S. scabiei* 87.22 |
| txtA_BsrGI_R | CACTGTACAGCTGCAGTGGGAGGTCCTGGACGCGGAG | |
| txtB_NdeI_F | CACCATATGTCCATGCTGCCGCCGGGGCGAAG | |
| txtB_BsrGI_R | CACTGTACAGCGGCCGTGGTGAGAAGGCCCGCCGTTC | |

**Table S3 – Annealing oligonucleotides – Cloning into pBP (EcoFlex)**

| Oligo ID | Sequence (5'-3') |
|---|---|
| SP44_A | TAGGTCTCACTATTGTTCACATTCGAACCGTCTCTGCTTTGACAACATGCTG |
| SP44_B | TGCGGTGTTGTAAAGTCTGGTGTAGTACAGAGACCCATG |
| SP44_C | CAGAGACGGTTCGAATGTGAACAATAGTGAGACC |
| SP44_D | GGTCTCTGTACTACACCAGACTTTACAACACCGCACAGCATGTTGTCAAAG |

**Table S4 – Sequencing primers**

| Oligo ID | Sequence (5'-3') | Template |
|---|---|---|
| hemD_seq | AACCCCACCAGCCCGAC | pSJM1147 |
| txtA_seq1 | GTCACCGGCGCCTCAC | pSJM1356 |
| txtA_seq2 | GCTCTACGACGATCTGTACG | |
| txtA_seq3 | CGTGGGAAACCGGAGC | |
| txtA_seq4 | CGAAGCCTGTGTCGCGAC | |
| txtB_seq1 | CCAGCCTGGTGGGCAGC | pSJM1357 |
| txtB_seq2 | CTCGTCGCCTATGTCGTG | |
| txtB_seq3 | GACGCGCAGACACTGAC | |
| txtB_seq4 | GTGCCGCTCTCGCAC | |

### DNA sequences

The 4-bp overhang for golden gate assembly between each part is as followed.
CTAT – Promoter – GTAC – Ribozyme – TTAA – RBS – CATA – Gene (mScarlet-I or sfGFP) – TCGA – Terminator – TGTT

**Table S5 – Promoter, RBS and RiboJ parts (unmodified SP12-SP44 also prepared)**

| Promoter | Activity[a] | SD[a] | Sequence[b] |
|---|---|---|---|
| P15/kasOp* | 100.00 | 0.55 | TGTTCACATTCGAACAGTCTCTGCTTTGACAACATGCTGTGCGGTGTTGTAAAGTCGTGGCCAGTAC |
| SP12a | 11.26 | 1.79 | TGTTCACATTCGAACAGTCTCTGCTTTGACAGGTCCATACACGCGCTTGTAAAGTCGTGGCCAGTAC |
| SP15a | 16.89 | 1.20 | TGTTCACATTCGAACAGTCTCTGCTTTGACACCTACGTGACACATCTTGTAAAGTCGTGGCCAGTAC |
| SP19a | 23.44 | 2.47 | TGTTCACATTCGAACAGTCTCTGCTTTGACACGACCCACTTTCGCTGTGTAAAGTCGTGGCCAGTAC |

| | | | |
|---|---|---|---|
| SP23a | 50.75 | 7.16 | TGTTCACATTCGAACAGTCTCTGCTTTGACAACATGCTGTGCGGTGTT GTAAAGTCCTGCTAAGTAC |
| SP25a | 58.99 | 0.72 | TGTTCACATTCGAACAGTCTCTGCTTTGACAGAGGTAGGCACGCTCA TGTAAAGTCGTGGCCAGTAC |
| SP30a | 75.14 | 5.66 | TGTTCACATTCGAACAGTCTCTGCTTTGACATCGTGTGGCGCTTGGG TGTAAAGTCGTGGCCAGTAC |
| SP40a | 116.55 | 5.21 | TGTTCACATTCGAACAGTCTCTGCTTTGACAACATGCTGTGCGGTGTT GTAAAGTCCCGTAGAGTAC |
| SP43a | 154.50 | 7.48 | TGTTCACATTCGAACAGTCTCTGCTTTGACACGGACAAGCGCTATGG TGTAAAGTCGTGGCCAGTAC |
| SP44a | 187.45 | 8.64 | TGTTCACATTCGAACAGTCTCTGCTTTGACAACATGCTGTGCGGTGTT GTAAAGTCTGGTGTAGTAC |
| *ermEp** | 5.09 | 0.19 | GGTACCAGCCCGACCCGAGCACGCGCCGGCACGCCTGGTCGATGT CGGACCGGAGTTCGAGGTACGCGG CTTGCAGGTCCAGGAAGGGGACGTCCATGCGAGTGTCCGTTCGAGT GGCGGCTTGCGCCCGATGCTAGTCGCGGTTGATCGGCGATCGCAG GTGCACGCGGTCGATCTTGACGGCTGGCGAGAGGTGCGGGGAGGA TCTGACCGACGCGGTCCACACGTGGCACCGCGATGCTGTTGTGGGC ACAATCGTGCCGGTTGGTAGGATCCGTAC |
| RBS | | | |
| R15 | 100 | 10.99 | TCTAAGTAAGGAGTGTCCATA |
| SR4 | 8.29 | 0.60 | GAACTTAAAGGAGTGTCCATA |
| SR7 | 18.11 | 0.67 | GCTACCCGTAGAGTGTCCATA |
| SR9 | 39.00 | 4.17 | CAAAAATCGTGAGTGTCCATA |
| SR12 | 59.27 | 1.32 | CGCGCCTAAGGAGTGTCCATA |
| SR18 | 87.10 | 0.86 | TCTCCTCAAGGAGTGTCCATA |
| SR24 | 109.34 | 1.84 | GCGTGTGAAGGAGTGTCCATA |
| SR28 | 130.30 | 5.83 | CATGCCATTGGAGTGTCCATA |
| SR35 | 156.90 | 1.92 | TAGCCTTAAGGAGTGTCCATA |
| SR39 | 174.23 | 11.22 | TTCGAGTAAGGAGTGTCCATA |
| SR40 | 183.52 | 22.78 | CTTGTCAAAGGAGTGTCCATA |
| Insulator | | | |
| RiboJ | NA | NA | AGCTGTCACCGGATGTGCTTTCCGGTCTGATGAGTCCGTGAGGACG AAACAGCCTCTACAAATAATTTTGTTTAA |

[a]Data obtained from Bai et al, PNAS 2015.
[b]Single base mutated (C>A) to remove internal BsmBI site.
[c]Ribozyme sequence obtained from Nielsen et al, Science 2016.

**sfGFP**

- Codon optimized for *S. venezuelae* ATCC 10712 (in pBP-ORF)
- Sub-cloned with NdeI/BamHI or EcoFlex assembly

GGTCTCACAT**ATG**TCCAAGGGCGAGGAGCTGTTCACCGGCGTCGTCCCGATCCTGGTCGAGCTGGACG
GCGACGTGAACGGCCACAAGTTCTCCGTCCGCGGCGAGGGCGAGGGCGACGCCACCAACGGCAAGCTG
ACCCTGAAGTTCATCTGCACCACCGGCAAGCTCCCGGTCCCGTGGCCGACCCTGGTCACCACCCTGAC
CTACGGCGTCCAGTGCTTCTCCCGCTACCCGGACCACATGAAGCGCCACGACTTCTTCAAGTCCGCCA
TGCCCGAGGGCTACGTCCAGGAGCGGACCATCTCCTTCAAGGACGACGGCACCTACAAGACCCGCGCC
GAGGTCAAGTTCGAGGGCGACACCCTGGTCAACCGCATCGAGCTGAAGGGCATCGACTTCAAGGAGGA
CGGCAACATCCTGGGCCACAAGCTCGAGTACAACTTCAACTCCCACAACGTCTACATCACCGCCGACA
AGCAGAAGAACGGCATCAAGGCCAACTTCAAGATCCGCCACAACGTCGAGGACGGCAGCGTCCAGCTG
GCCGACCACTACCAGCAGAACACCCCGATCGGCGACGGCCCGGTCCTGCTGCCGGACAACCACTACCT
GTCCACCCAGTCCGTCCTGTCCAAGGACCCGAACGAGAAGCGCGACCACATGGTCCTGCTCGAGTTCG
TCACCGCCGCCGGCATCACCCACGGCATGGACGAGCTGTACAAG**TGA**GGATCCTCGAAGAGACC

**mScarlet-I**

- Codon optimized for *S. venezuelae* ATCC 10712 (in pBP-ORF)
- Flag- and FlAsH-EDT$_2$ tags and dBrocolli mRNA aptamer
- Sub-cloned from gBlock fragment into pTU1-SP44-PET using NdeI and BglII/BamHI

CAT**ATG**GTGTCCAAGGGCGAGGCCGTCATCAAGGAGTTCATGCGCTTCAAGGTCCACATGGAGGGCTC
CATGAACGGCCACGAGTTCGAGATCGAGGGCGAGGGCGAGGGCCGCCCGTACGAGGGCACCCAGACCG
CCAAGCTGAAGGTCACCAAGGGCGGCCCGCTGCCGTTCTCCTGGGACATCCTGTCCCCGCAGTTCATG
TACGGCTCCCGCGCCTTCATCAAGCACCCGGCCGACATCCCGGACTACTACAAGCAGTCCTTCCCGGA
GGGCTTCAAGTGGGAGCGCGTCATGAACTTCGAGGACGGCGGCGCCGTCACCGTCACCCAGGACACCT
CCCTGGAGGACGGCACCCTGATCTACAAGGTCAAGCTGCGCGGCACCAACTTCCCGCCGGACGGCCCG
GTCATGCAGAAGAAGACCATGGGCTGGGAGGCCTCCACCGAGCGCCTGTACCCGGAGGACGGCGTCCT
GAAGGGCGACATCAAGATGGCCCTGCGCCTGAAGGACGGCGGCCGCTACCTGGCCGACTTCAAGACCA
CCTACAAGGCCAAGAAGCCGGTCCAGATGCCGGGCGCCTACAACGTCGACCGCAAGCTGGACATCACC
TCCCACAACGAGGACTACACCGTCGTCGAGCAGTACGAGCGCTCCGAGGGCCGCCACTCCACCGGCGG
CATGGACGAGCTGTACAAGGGATCCGACTACAAGGACGACGACGACAAGTGCTGCCCGGGCTGCTGC**T**
**AA**TTGCCATGTGTATGTGGGAGACGGTCGGGTCCAGATATTCGTATCTGTCGAGTAGAGTGTGGGCTC
CCACATACTCTGATGATCCTTCGGGATCATTCATGGCAAAGATCT

**mVenus-I**

- Codon optimized for *S. venezuelae* ATCC 10712 (in pBP-ORF)
- Flag- and FlAsH-tags
- Sub-cloned from gBlock fragment into pTU1-SP44-PET-sfGFP-Bba_B0015 using NdeI and BglII/BamHI

CAT**ATG**GTGTCCAAGGGCGAGGAGCTGTTCACCGGCGTCGTCCCGATCCTGGTCGAGCTGGACGGCGA
CGTCAACGGCCACAAGTTCTCCGTCTCCGGCGAGGGCGAGGGCGACGCCACCTACGGCAAGCTGACCC
TGAAGCTGATCTGCACCACCGGCAAGCTGCCGGTCCCGTGGCCGACCCTGGTCACCACCCTGGGCTAC
GGCCTGCAGTGCTTCGCCCGCTACCCGGACCACATGAAGCAGCACGACTTCTTCAAGTCCGCCATGCC
GGAGGGCTACGTCCAGGAGCGCACCATCTTCTTCAAGGACGACGGCAACTACAAGACCCGCGCCGAGG
TCAAGTTCGAGGGCGACACCCTGGTCAACCGCATCGAGCTGAAGGGCATCGACTTCAAGGAGGACGGC
AACATCCTGGGCCACAAGCTGGAGTACAACTACAACTCCCACAACGTCTACATCACCGCCGACAAGCA
GAAGAACGGCATCAAGGCCAACTTCAAGATCCGCCACAACATCGAGGACGGCGGCGTCCAGCTGGCCG
ACCACTACCAGCAGAACACCCCGATCGGCGACGGCCCGGTCCTGCTGCCGGACAACCACTACCTGTCC
TACCAGTCCAAGCTGTCCAAGGACCCGAACGAGAAGCGCGACCACATGGTCCTGCTGGAGTTCGTCAC
CGCCGCCGGCATCACCCTGGGCATGGACGAGCTGTACAAGAGATCTGACTACAAGGACGACGACGACA
AGTGCTGCCCGGGCTGCTGC**TAA**ACAATAACTGAATAGGGGATCC

**β-D-glucuronidase (GUS) (Escherichia coli str. K-12 substr. MG1655)**

- Codon optimized for *S. venezuelae* ATCC 10712
- Sub-cloned with NdeI/BamHI or EcoFlex assembly

GGTCTCACAT**ATG**CTGCGCCCGGTCGAAACCCCGACCCGCGAGATCAAGAAGCTGGACGGCCTGTGGG
CCTTCTCCCTGGACCGCGAGAACTGCGGCATCGACCAGCGCTGGTGGGAGTCCGCCCTGCAGGAGTCC
CGCGCCATCGCCGTCCCGGGCTCCTTCAACGACCAGTTCGCCGACGCCGACATCCGCAACTACGCCGG
CAACGTCTGGTACCAGCGCGAGGTCTTCATCCCGAAGGGCTGGGCCGGCCAGCGCATCGTCCTGCGCT
TCGACGCCGTCACCCACTACGGCAAGGTCTGGGTCAACAACCAGGAGGTCATGGAGCACCAGGGCGGC
TACACCCCGTTCGAGGCCGACGTCACCCCGTACGTCATCGCCGGCAAGTCCGTCCGCATCACCGTCTG
CGTCAACAACGAGCTGAACTGGCAGACCATCCCGCCGGGCATGGTCATCACCGACGAGAACGGCAAGA
AGAAGCAGTCCTACTTCCACGACTTCTTCAACTACGCCGGCATCCACCGCTCCGTCATGCTGTACACC
ACCCCGAACACCTGGGTCGACGACATCACCGTCGTCACCCACGTCGCCCAGGACTGCAACCACGCCTC
CGTCGACTGGCAGGTCGTCGCCAACGGCGACGTGTCCGTCGAGCTGCGCGACGCCGACCAGCAGGTCG
TCGCCACCGGCCAGGGCACCTCCGGCACCCTGCAGGTCGTCAACCCGCACCTGTGGCAGCCGGGCGAG
GGCTACCTGTACGAGCTGTGCGTCACCGCCAAGTCCCAGACCGAGTGCGACATCTACCCGCTGCGCGT
CGGCATCCGCTCCGTCGCCGTCAAGGGCGAGCAGTTCCTGATCAACCACAAGCCGTTCTACTTCACCG
GCTTCGGCCGCCACGAGGACGCCGACCTGCGCGGCAAGGGCTTCGACAACGTCCTGATGGTCCACGAC
CACGCCCTGATGGACTGGATCGGCGCCAACTCCTACCGCACCTCCCACTACCCGTACGCCGAGGAGAT
GCTGGACTGGGCCGACGAGCACGGCATCGTCGTCATCGACGAAACCGCCGCCGTCGGCTTCAACCTGT
CCCTGGGCATCGGCTTCGAGGCCGGCAACAAGCCGAAGGAGCTGTACTCCGAGGAGGCCGTCAACGGC
GAAACCCAGCAGGCCCACCTCCAGGCCATCAAGGAGCTGATCGCCCGCGACAAGAACCACCCGTCCGT
CGTCATGTGGTCCATCGCCAACGAGCCGGACACCCGCCCGCAGGGCGCCCGCGAGTACTTCGCCCCGC
TGGCCGAGGCCACCCGCAAGCTGGACCCGACCCGCCCGATCACGTGCGTCAACGTCATGTTCTGCGAC

GCCCACACCGACACCATCTCCGACCTGTTCGACGTCCTGTGCCTGAACCGCTACTACGGCTGGTACGT
CCAGTCCGGCGACCTGGAAACCGCCGAGAAGGTCCTGGAGAAGGAGCTGCTGGCCTGGCAGGAGAAGC
TGCACCAGCCGATCATCATCACCGAGTACGGCGTCGACACCCTGGCCGGCCTGCACTCCATGTACACC
GACATGTGGTCCGAGGAGTACCAGTGCGCCTGGCTGGACATGTACCACCGCGTCTTCGACCGCGTGTC
CGCCGTCGTCGGCGAGCAGGTCTGGAACTTCGCCGACTTCGCCACCTCCCAGGGCATCCTGCGCGTCG
GCGGCAACAAGAAGGGCATCTTCACCCGCGACCGCAAGCCGAAGTCCGCCGCCTTCCTGCTGCAGAAG
CGCTGGACCGGCATGAACTTCGGCGAGAAGCCGCAGCAGGGCGGCAAGCAG**TGA**GGATCCTCGAAGAG
ACC

**OxyD (*S. rimosus* DSM-40260)**

- Codon optimized for *S. venezuelae* ATCC 10712
- Sub-cloned with NdeI/BamHI or EcoFlex assembly
- No stop codon

GGTCTCACAT**ATG**TGCGGCATCGCCGGCTGGATCGACTTCGAGCGCAACCTCGCGCAGGAGCGCGCCA
CCGCCTGGGCCATGACCGACACCATGGCCTGCCGCGGCCCGGACGACGCCGGCCTCTGGACCGGCGGC
CACGCCGCGCTCGGCCACCGCCGCCTGGCCGTCATCGACCCGGCGCACGGCCGCCAGCCGATGCACTC
GACCCTGCCGGACGGCACCTCGCACGTCATCACCTTCTCCGGCGAGATCTACAACTTCCGCGAGCTGC
GCGTCGAGCTGGAGTCCCAGGGCCACCGCTTCCGCACGCACTGCGACACCGAGGTCGTCCTCCACGGC
TACACCCGCTGGGGCCGCGAGCTGGTCGACCGCCTCAACGGCATGTACGCCTTCGCCGTCTGGGACGA
GGCCCGCCAGGAGCTGCTCCTCGTCCGCGACCGCATGGGCGTCAAGCCGCTCTACTACCACCCGACCG
CCACCGGCGTCCTCTTCGGCAGCGAGCCGAAGGCCGTCCTCGCGCACCCGTCGCTCCGCCGCCGCGTC
ACCGCCGAGGGCCTCTGCGAGGTCCTCGACATGGTCAAGACCCCGGGCCGCACCGTGTTCTCCGGCAT
GCGCGAGGTGCTCCCGGGCGAGATGGTCACCGTCGGCCGCTCCGGCGTCGCCCGCCGGCGCTACTGGA
CGCTGCAGGCCCGCGAGCACACCGACGACCTCGAAACCACGATCGCGACCGTCCGCGGCCTCCTCACC
GACACCGTCCGCCGCCAGCTCGTCAGCGACGTCCCGCTCTGCACCCTCCTCTCCGGCGGCCTCGACTC
CTCCGCCGTCACCGCGCTCGCCGCGCGCGCCGGCGACGGCCCGGTCCGCACCTTCTCGGTCGACTTCA
GCGGCGCGGGCACCCGCTTCCAGCCGGACGCGGTCCGCGGCAACACGGACGCCCCGTACGTCCAGGAG
ATGGTCCGCCACGTCGCGGCCGACCACACCGAGGTGGTGCTCGACTCCGCCGACCTCGCGGCGCCCGA
GGTGCGCGCCGCGGTCCTCGGCGCGACCGACCTGCCGCCGGCCTTCTGGGGCGACATGTGGCCGTCGC
TCTACCTCTTCTTCCGCCAGGTCCGCCAGCACTGCACCGTCGCGCTGTCCGGCGAGGCGGCCGACGAG
CTGTTCGGCGGCTACCGCTGGTTCCACCGGACCGCCGCCATCGACGCCGGCACGTTCCCGTGGCTCAC
CGCCGGCAGCGCCCGCTACTTCGGCGGCCGGGGCCTCTTCGACCGCAAGCTCCTCGACAAGCTGGACC
TGCCGGGCTACCAGCGCGACCGCTACGCCGAGGCGCGCAAGGAGGTCCCGGTCCTGCCCGGCGAGGAC
GCCCGCGAGGCGGAGCTGCGGCGGGTCACCTACCTCAACCTCACGCGCTTCGTGCAGACCCTCCTGGA
CCGCAAGGACCGCATGTCCATGGCCACCGGCCTCGAGGTCCGCGTGCCGTTCTGCGACCACCGCCTCG
TCGACTACGTGTTCAACGTCCCGTGGGCCATGAAGTCCTTCGACGGCCGCGAGAAGTCCCTCCTCCGC
GCCGCCGTCCGGGACCTGCTGCCGGAGTCCGTCGTCACCCGCGTCAAGACGCCGTACCCGGCCACGCA
GGACCCGGTCTACGAGCGCCTGCTCCGGGACGAGCTGGCCGCGCTCCTCGCCGACTCGCAGGCGCCGG
TCCGCGAGCTGCTGGACCTCGGCCGGGCGCGCGACCTGCTCCGCCGGCCGGTCGGCGCCGTCAGCCAG
CCGTACGACCGCGGCTCCCTCGAGCTGGTGCTCTGGATGAACACCTGGCTCGCGGAGTACGGCGTGTC
GCTCGAGCTGGGATCCTCGAAGAGACC
